# Blue-green opponency and trichromatic vision in the greenhouse whitefly (*Trialeurodes vaporariorum*)

**DOI:** 10.1101/341610

**Authors:** Niklas Stukenberg, Hans-Michael Poehling

**Affiliations:** Leibniz-Universität Hannover, Institute of Horticultural Production Systems, Section Phytomedicine, Herrenhäuser Str. 2, 30419 Hannover, Germany

**Keywords:** Wavelength-specific behaviour, visual behaviour, opponent chromatic mechanism, colour vision, colour choice model, LEDs

## Abstract

Visual orientation in the greenhouse whitefly (*Trialeurodes vaporariorum* Westwood, Hemiptera: Aleyrodidae) is the result of ‘wavelength-specific behaviours’. Green-yellow elicits ‘settling behaviour’ while ultraviolet (UV) radiation initiates ‘migratory behaviour’. Electroretinograms of the photoreceptors’ spectral efficiency showed peaks in the green and the UV range and whitefly vision was said to be dichromatic.

In order to study the visual behaviour of *T. vaporariorum*, nineteen narrow-bandwidth LEDs covering the UV-A and visible range were used in combination with light scattering acrylic glass screens in a small-scale choice arena under greenhouse conditions. Multiple-choice and dual-choice assays were performed, resulting in LED-based behavioural action spectra of settling (green) and migratory behaviour (UV). A potential inhibitory blue-green chromatic mechanism was studied by combining yellow with different blueish LEDs. Intensity dependencies were illustrated by changing LED intensities.

Regarding the ‘settling response’, highest attraction was achieved by a green LED with a centroid wavelength of 550 nm, while a blue LED with 469 nm proved to be most inhibitory. Behaviour was distinctly intensity dependent. ‘Migratory behaviour’ was elicited the most by the UV LED with the shortest available wavelength of 373 nm. The results clearly prove the presence of a green and a yet undescribed blue sensitive photoreceptor and a blue-green opponent mechanism. Furthermore, empirical colour choice models were built and receptor peaks were estimated around 510 - 520 nm (green), 480 - 490 nm (blue) and 340 - 370 nm (UV). Consequently, *Trialeurodes vaporariorum* possesses a trichromatic receptor setup.

**Summary statement:** LED based choice experiments and empirical colour choice models reveal a yet undescribed blue sensitive photoreceptor and an inhibitory interaction with a green sensitive receptor.

## Introduction

Visual orientation is crucial for initial host plant detection and migration in the greenhouse whitefly (*Trialeurodes vaporariorum* Westwood, Hemiptera: Aleyrodidae), a worldwide occurring horticultural pest in greenhouses (Byrne, 1991). Two different behavioural patterns, so called ‘wavelength-specific behaviours’, were identified in *T. vaporariorum*. Orientation to host plants is guided by a ‘settling’ behaviour which is elicited by green-yellow light while ultraviolet (UV) radiation is responsible for a pattern which can be broadly defined as ‘migratory behaviour’ (Coombe, 1981; 1982).

Those ‘wavelength-specific behaviours’ are generally defined as innate colour-sensitive behavioural responses to different wavelength bands which cannot be modified by experience or learning. On a basic level they enable insects to find and discriminate targets by their specific patterns of reflected light (Kelber and Osorio, 2010). In herbivorous insects the green-yellow range is commonly used for host plant detection (Prokopy and Owens, 1983). UV radiation is generally known to be involved in spatial orientation, flight activity, and dispersal in a variety of insects (Briscoe and Chittka, 2001).

The physiological basis for the visual perception of light are the photoreceptor cells in the insects’ compound eyes containing the visual pigments. The absorption spectrum of visual pigments can be expressed by its sensitivity function which can be described using template formulas (Govardovskii et al., 2000; Kelber et al., 2003). According to the principle of univariance, a single photoreceptor is colour-blind because wavelength and intensity-dependent stimulation are confounded. The receptor screens a certain wavelength range but the same signal can be elicited by low intensity light at the sensitivity peak wavelength or by high intensity light further away from peak sensitivity (Skorupski and Chittka, 2011; Naka and Rushton, 1966).

‘Wavelength-specific behaviour’ can be based on the output of a single photoreceptor and achromatic, i.e. brightness-related, processing. Furthermore, it can be the result of colour opponency which is a chromatic mechanism in which the outputs of several photoreceptors are compared by antagonistic neuronal processing. Colour opponency is a prerequisite of colour vision defined as the ability to detect spectral variations in the light independent of their intensity (Skorupski and Chittka, 2011; Kemp et al., 2015; Kelber and Osorio, 2010; Kelber et al., 2003).

Many studies indicate that for herbivorous insects such as aphids, the ‘settling’ behaviour is controlled by such an inhibitory interaction of two overlapping photoreceptors sensitive for blue and green light. In this so called ‘opponent mechanism’ or ‘blue-green opponency’ the signal from the blue receptor inhibits the signal from the green receptor eliciting ‘settling’ (Döring and Chittka, 2007; Döring, 2014; Döring and Röhrig, 2016; Döring et al., 2009). This mechanism facilitates to extract a constant chromatic signal that detects reflected long-wavelength light (green-yellow) associated with host plants and discriminates it from short- or broad-wavelength light independent from illumination intensity. It also results in a shift of the behavioural action spectrum to the longer wavelength range as compared to the underlying photoreceptor sensitivity and a more specific and narrow tuning in to the relevant green wavelength range. An apparent shortcoming of this dichromatic mechanism is the common preference of many herbivorous insects for yellow instead of green which can be explained by higher reflection in the relevant green range resulting in higher relative input to the green receptor. Therefore, this simple chromatic mechanism, which should be independent of light intensity, is influenced by brightness in terms of changing blue and green photoreceptor excitation ratios. Thereby, it may be that the whole mechanism lies on a mixed achromatic and chromatic axis (Döring and Chittka, 2007; Kelber and Osorio, 2010; Skorupski and Chittka, 2011).

Similar to aphids and other herbivorous insects, *Trialeurodes vaporariorum* shows a clear preference for yellow-reflecting objects. At an early stage, Moericke et al. (1966) identified a ‘fall reflex’ consistently elicited above yellow surfaces independent of the intensity of the reflected colour and suggested some form of ‘wavelength-specific behaviour’ or colour vision. This preference for yellow was later confirmed in behavioural studies with coloured surfaces, and bright yellow with little to no reflectance in the violet-blue spectrum was identified as being most attractive compared to darker or less saturated yellow. Violet-blue proved to be not attractive and it even inhibits the attraction towards yellow. Moreover, it was shown that highly reflected intensities in the green-yellow range contribute positively to their attractiveness (Vaishampayan et al., 1975; Affeldt et al., 1983; Webb et al., 1985). All these results with coloured surfaces have contributed to the development and use of yellow sticky traps for monitoring and control of whiteflies in horticultural greenhouse crops (Böckmann et al., 2015; Gillespie and Quiring, 1987).

In a behavioural study with monochromatic light of controlled intensities MacDowall (1972) determined the spectral efficiency function for a wavelength pattern from blue to red. The revealed action spectrum peaked at 550 nm and corresponded with the reflection spectrum of a tobacco leaf. Coombe (1981) extensively investigated the visual behaviour using monochromatic light in a ‘settling’ paradigm and a ‘phototactic’ paradigm. An action spectrum for the ‘settling response’ was generated based on spectral sensitivity which peaked at 550 nm and had a second peak in the UV range at 350 nm. Based on intensity response functions and different methods for the determination of ‘settling’ it was concluded that *T. vaporariorum* exhibits ‘wavelength-specific behaviour’. In the phototactic paradigm it could be shown that two different antagonistic behavioural patterns are elicited by 400 nm (UV) and 550 nm (green) which do not interact with each other. In a follow-up study (Coombe, 1982), it was further revealed that UV elicits a variety of responses associated with migratory behaviour, such as take-off behaviour and maintenance of flight. For example, increased walking activity and takeoff rates were observed under 400 nm light and UV was preferred over green light but only during flight activity. In accordance with that, it is reported from many applied studies that whiteflies show less flight activity in UV-deficient environments leading to a general avoidance of such conditions (Gulidov and Poehling, 2013; Kumar and Poehling, 2006; Antignus et al., 2001).

For aphids, clear physiological evidence of a trichromatic receptor setup involving UV-sensitive photoreceptors exists (Kirchner et al., 2005). In contrast, trichromacy has not been confirmed in *Trialeurodes vaporariorum*. Mellor et al. (1997) investigated the physiological properties of the compound eye of *T. vaporariorum* and determined its spectral efficiency using the electroretinogram (ERG) technique. Efficiency peaks were identified in the green-yellow region (520 nm) and in the UV region (340 nm). Furthermore, the eye is divided in a dorsal part with 54-55 ommatidia and a ventral part containing 29-31 ommatidia. The dorsal region was thereby more sensitive to UV. Based on these results the visual system was concluded to be dichromatic.

New insights could be achieved by Stukenberg et al. (2015) using choice experiments with narrow bandwidth light emitting diodes (LEDs). Green LED traps were preferred over yellow sticky traps but this attraction was supressed when simultaneously combined with blue LEDs. This is the first clear indication that a yet undetected blue photoreceptor close to a green receptor and an inhibitory chromatic interaction between both might be present in the greenhouse whitefly. A moderate attractiveness towards UV could also be shown and it seemed to have an enhancing or synergistic effect on the attractiveness of green light as the combination of UV and green LEDs was more attractive than green alone, especially under night-time conditions. In a recent study, yellow rollertraps with reduced translucency were more attractive than those with common translucency. The authors determined the spectral properties of the traps and explained the results on the basis of the potential blue-green opponency. The brighter reflection in the green-yellow range and the low transmission of blue light had a greater influence on the opponent mechanism, resulting in higher attraction (Sampson et al., 2018).

Considering the referred studies it is quite likely that *T. vaporariorum* exhibits blue-green opponency and possesses a trichromatic receptor setup. Nevertheless, a clear proof and a detailed characterisation of the mechanism which connects behavioural data with potential photoreceptor sensitivities is still missing. LEDs are a very useful tool to study insects’ visual behaviour since wavelengths and intensities can be individually adjusted and combined (Tokushima et al., 2016; Booth et al., 2004). In this study, we explored the visual behaviour and wavelength discrimination ability of *T. vaporariorum* using a fine-tuned selection of LEDs ranging from UV to red. Behavioural action spectra were generated under semi-natural greenhouse conditions, thereby taking changing ambient light conditions into account. We further investigated and characterized in detail the potential blue photoreceptor and the blue-green chromatic mechanism by LED mixing experiments. From the data, we built simple empirical colour choice models which explain the choice behaviour and enable approximate estimation of the spectral location of photoreceptors.

## Material and Methods

### Experimental LED trap screens

In order to study the visual behaviour of *T. vaporariorum*, nineteen individual high-power (HP) light emitting diodes (LEDs) covering the UV-A and visible spectra were selected (Table 1, Fig.1). LEDs underlie limitations concerning wavelength availability and homogeneity of bandwidths and intensities and show variations among equally coloured LEDs. Criteria for the selection were the fine-tuned fitting to the spectral regions of interest, narrow bandwidths, and sufficient spectral distances and intensities. In the selection process, spectra of various HP LEDs were recorded with the spectrometer Avaspec 2048-2 (Avantes, Apeldoorn, The Netherlands).

**Table 1.**
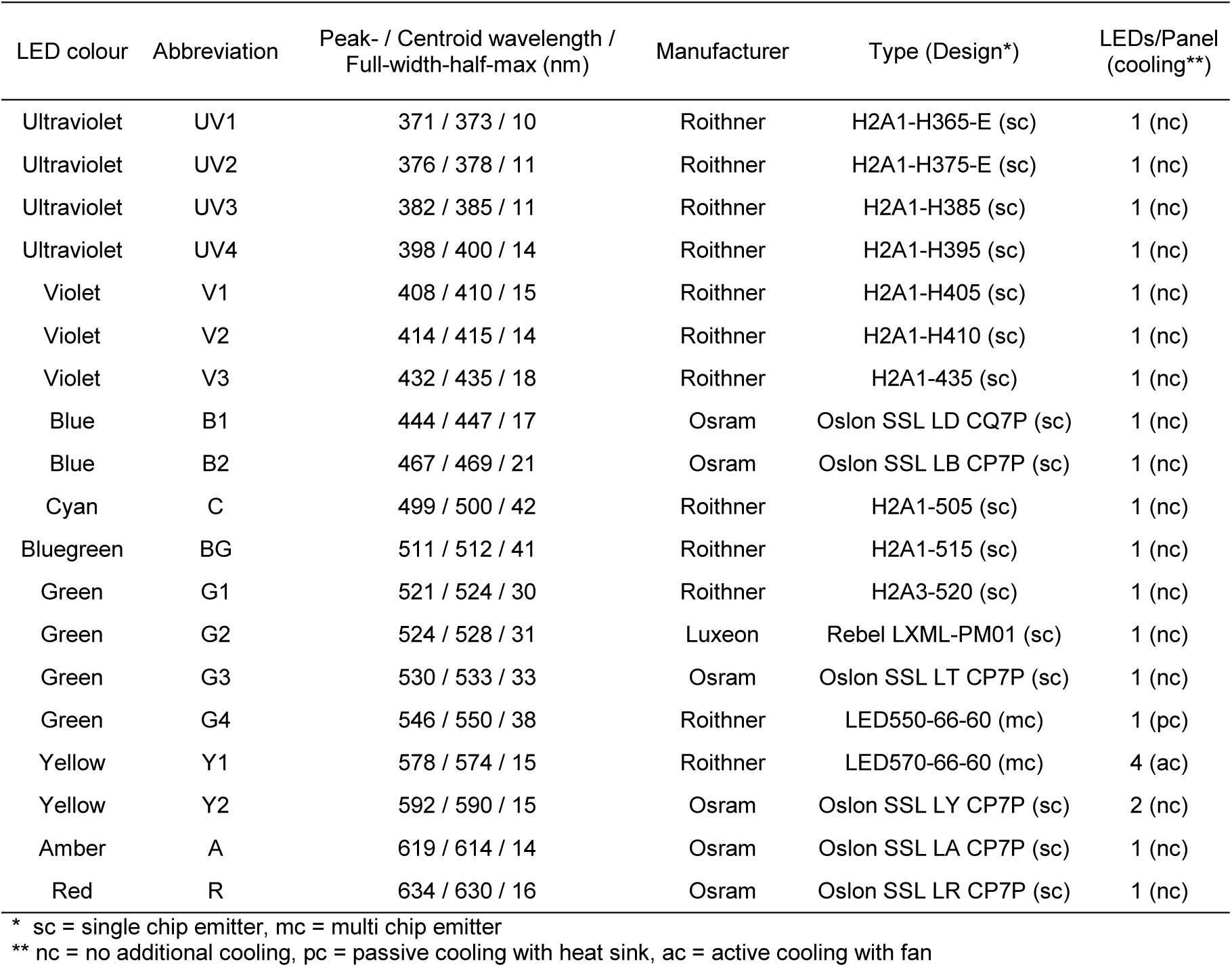
Specifications of high-power LEDs and constructed LED panels used in the experiments.

**Fig. 1.**
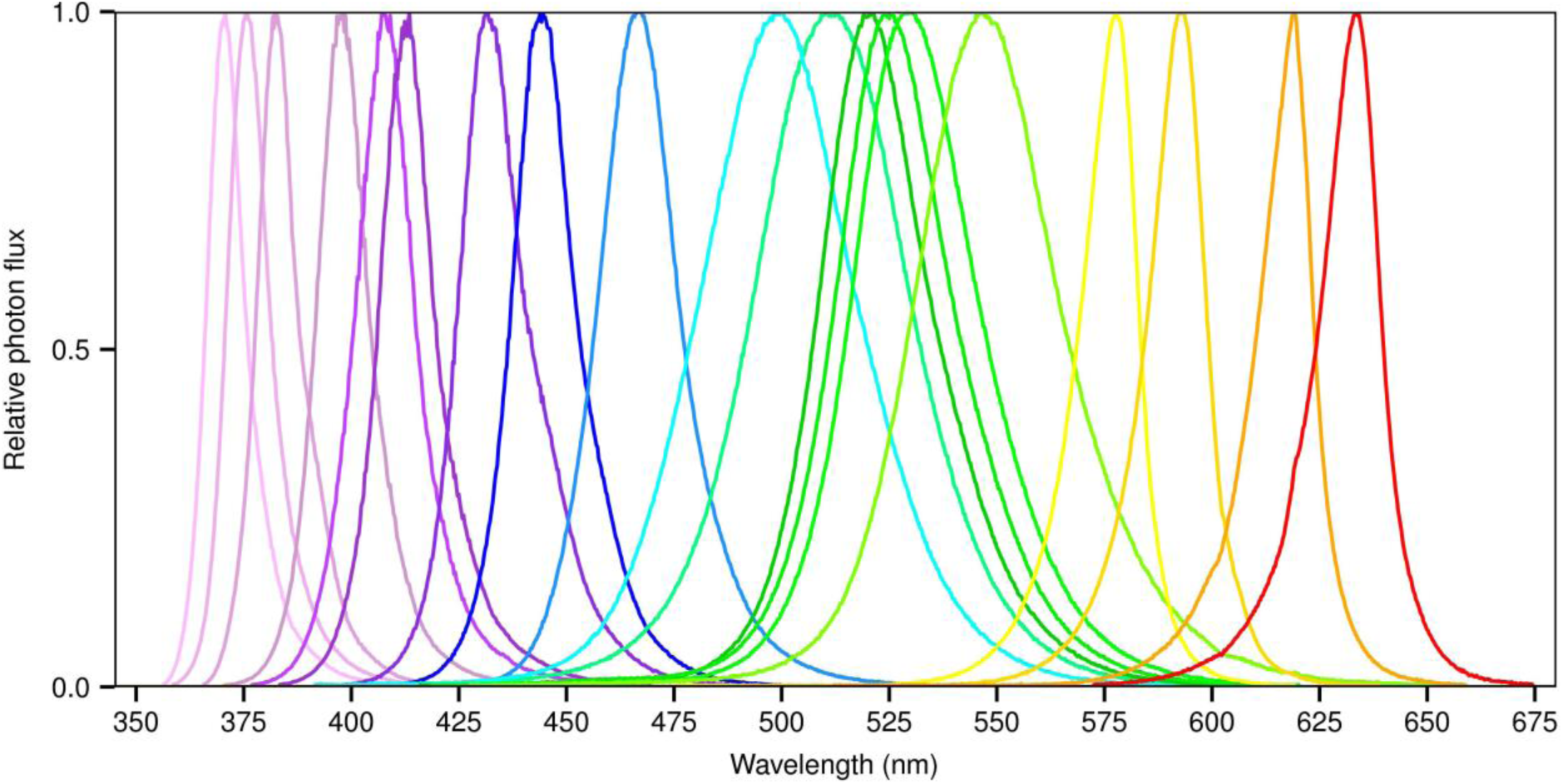
Spectra of high-power LEDs used. Data refer to LED specifications given in Table 1, in spectral order.

LEDs of each colour were attached to aluminium-panels (100 × 100 × 1 mm). To obtain sufficient intensities for yellow LEDs, two or four LEDs had to be used. Most HP LEDs were common single chip emitters but for chartreuse green and yellow specific multichip emitters had to be used (Table 1). They required additional cooling by heat sinks (Fischer Elektronik GmbH & Co. KG, Lüdenscheid, Germany) or even active cooling with a fan (LED cooling module, LA001-011A9DDN, Sunonwealth Electric Machine Industry Co., Ltd, Kaohsiung City, Taiwan).

As LED traps, boxes (0.1 × 0.1 × 0.13 m) were constructed out of grey PVC (4 mm) to insert the LED panels on the backside via grooves in the side walls. The front side of the box was closed by transparent a opal acrylic glass plate (100 × 100 × 3 mm, PLEXIGLAS^®^ LED 0M200 SC, Evonik Industries AG, Essen, Germany) which served as scatter screens (Fig. 2A). In addition, mirror film (PEARL GmbH, Buggingen, Germany) was used to laminate the insides of the boxes. For whitefly trapping, the screen was covered with transparent plastic film (PET) coated with insect glue (Temmen GmbH, Hattersheim, Germany), which was shown in preliminary tests to not influence the emitted spectra.

**Fig. 2.**
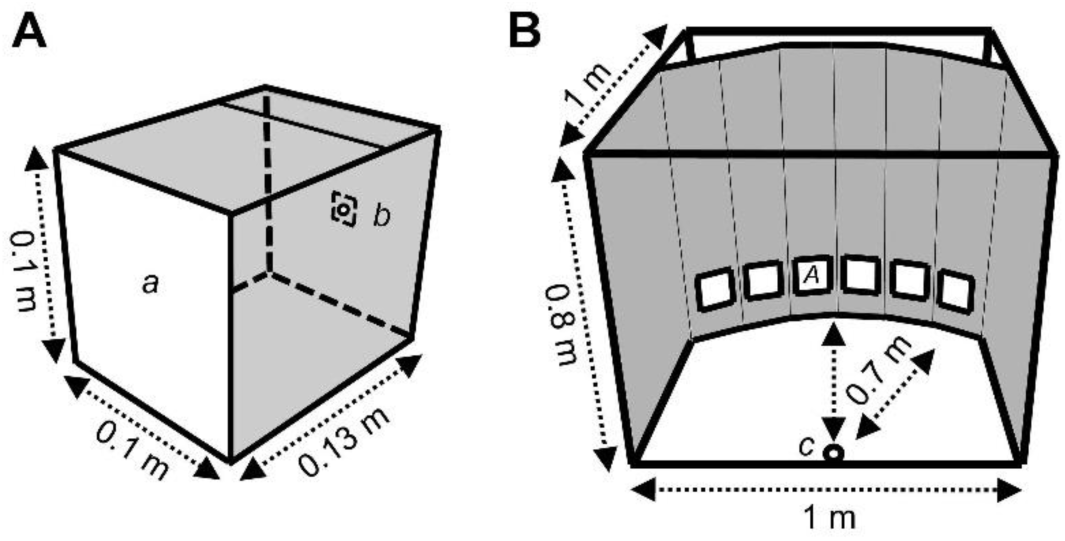
Schemes of LED trap screen and choice arena. (A) LED trap screen with acrylic glass screen front side (*a*) and LED panel backside (*b*). The inner side of the box was laminated with mirror film. (B) Choice arena with whitefly release point (*c*) and position of LED traps (*A*). The background was black and the bottom was black-brown.

For the operation and adjustment of intensities of each LED panel, a device with 16 LED drivers (Mini Jolly, TCI, Saronno, Italy) was constructed. The 16 separate channels could be dimmed (0-100%) by external control signals (0-10 V) which were provided by two USB analogue output modules (ME RedLab 3104, Meilhaus Electronic GmbH, Alling, Germany) in combination with a notebook and the software ProfiLab-Epert 4.0 (ABACOM, Ganderkesee, Germany).

Photon flux densities (µmol ^m-2^ s^-1^) of LEDs from the long-wave UV-A to red (UV3 - R, Table 1, Fig. 1) were measured and adjusted using the LI-250 A Light Meter with LI 190 Quantum Sensor (LI-COR Biosciences, Lincoln, NE, USA). As the sensor is only suitable to measure broadband photosynthetic active radiation (PAR, 400 – 700 nm), the sensor sensitivity data provided by LI-COR (starting at 385 nm) was included in the measurement of UV and violet LEDs (UV3 – V3, Table 1, Fig. 1). Extrapolation of the non-measurable parts of LED spectra below 385 nm had to be conducted. For the other two UV-A LEDs (UV1, UV2, Table 1, Fig. 1), the Almemo^®^ 2390-5 datalogger (Ahlborn Mess- und Regelungstechnik GmbH, Holzkirchen, Germany) in combination with a UV-A sensor (Type 2.5, Indium Sensor GmbH, Neuenhagen, Germany) were used. The intensities were indicated in W ^m-2^ and were converted to µmol m^-2^ s^-1^ using the LED spectra, Planck’s constant, and Avogadro’s number. The sensitivity data of the sensor was included as the sensor is matched for UV-A measurement in broadband sunlight. All measurements were conducted in darkness by placing the sensor directly on the centre of the LED screen surface.

### Whiteflies

Greenhouse whiteflies (*Trialeurodes vaporariorum*) were reared on tobacco plants (*Nicotiana tabacum* L. cv. ‘Xanthi’) in two gauze cages (0.75 × 0.5 × 0.8 m) at the Leibniz-Universität, Hannover, Institute of Horticultural Production Systems, Section Phytomedicine in Germany at 23 ± 3 °C. For each experimental trial, vital individuals were carefully collected with an aspirator from the underside of the top leaves into a snap-on lid glass vial (h × d = 50 × 30 mm) and immediately released into the experimental choice arena.

### LED choice arena

Choice experiments were conducted close to the whitefly rearing in the same greenhouse compartment. A gauze-covered flight cage (1 × 1 × 0.8 m, Fig. 2B) with a waterproof black-brown plywood bottom was placed on stands at a height of one meter. The foldable front side faced in northern direction and was equipped with an additional lockable circular opening (0.25 m diameter) enabling the releasing of whiteflies. A semicircular background made of carton sprayed with matt black acrylic paint (Dupli Color, Motip Dupli GmbH, Hassmersheim, Germany) was inserted into the cage at a distance of 0.7 m to the release point. The background was equipped with six square holes of 0.1 × 0.1 m at a height of 0.1 m and a distance of 0.05 m to each other. The LED trap screens could be optionally inserted from the backside by placing them on 0.1 m high wooden blocks (Fig. 2A,B). The cage backside was covered with gauze and black-silver reflective mulch film (Sunup Reflective Films, Oceanside, CA, USA). The cables for each LED panel were connected from the cage backside to the LED control placed under the cage.

The ambient solar radiation during the experiments was measured using a sensor for visible light (FLA 623 PS, Ahlborn Mess- und Regelungstechnik GmbH, Holzkirchen, Germany) and a UV-A sensor (300 – 400 nm, Type 2.5, Indium Sensor GmbH, Neuenhagen, Germany) placed next to the whitefly release point. Measurements were recorded at 20 second intervals with the Almemo^®^ 2590-4AS datalogger (Ahlborn Mess- und Regelungstechnik GmbH, Holzkirchen, Germany) which was also placed under the cage. Temperature was recorded with a Tinytag Plus 2 TGP-4500 datalogger (Gemini Data Loggers Ltd., Chichester, UK).

### Experimental overview and classification

According to literature, the behavioural response to the green-yellow range corresponds with ‘settling’ while the response to the UV-violet range is presumably related to ‘migratory behaviour’ (Coombe, 1981; 1982). The conducted experiments can be classified into wavelength dependence experiments characterized by the predominant main colours (green, blue, UV) and intensity experiments in the green-yellow range, resulting in four experimental blocks (Table 2). Wavelength dependence experiments on the ‘settling response’ are referred to as ‘Green response experiments’ (Block 1). Subsequently, ‘intensity dependencies’ in the green-yellow range were determined (Block 2). An inhibitory blue-green chromatic mechanism in the ‘settling response’ was studied by combining yellow LEDs of the same wavelength with blueish LEDs of different wavelengths, referred to as ‘Blue inhibition experiments’ (Block 3). Wavelength dependence experiments of the ‘migratory behaviour’ are referred to as ‘UV response experiments’ (Block 4).

**Table 2.** Experimental overview and design

| Block | Experimental | | LEDs ** | Intensity on trap screen<br>($\mu\text{mol m}^{-2} \text{s}^{-1}$) | Number of | | | Trial Duration<br>(h) *** | Dates |
| --- | --- | --- | --- | --- | --- | --- | --- | --- | --- |
|  | No. | Design * |  |  | Replicates | Days | Trials / day | Whitflies / trial |  |
| (1)<br>Green<br>response | 1 | m-c | B2, C, BG, G1, G2, G3 | 28 | 20 | 5 | 4 | 150 | 12-17/02/2015 |
|  | 2 | m-c | G3, G4, Y1, Y2, A, R | 28 | 20 | 5 | 4 | 150 | 06-11/02/2015 |
|  | 3 | m-c | G1, G2, G3, G4, Y1, Y2 | 28 | 20 | 5 | 4 | 200 | 20-25/02/2015 |
|  | 4 | m-c | B1, B2, C, BG, A, R | 28 | 10 | 2 | 5 | 150 | 09-10/04/2015 |
|  | 5 | d-c | G4 vs.<br>BG, G1, G2, G3, Y1, Y2, A | 28 | 10 | 10 | 7 | 150 | 02-16/03/2015 |
| (2)<br>Intensity<br>dependence | 6 | d-c | G4 at 50% intensity vs.<br>G1, G3, Y1, Y2 at 100% | 14 vs. 28 | 10 | 5 | 8 | 150 | 23-30/03/2015 |
|  | 7 | m-c | Y2 at different intensities<br>(100, 83, 66, 50, 33, 16%) | 54, 45, 36, 27, 18, 9 | 20 | 5 | 4 | 200 | 24-27/11/2015 |
|  | 8 | m-c | Y2 at equal intensities | 50 | 10 | 2 | 5 | 200 | 15-16/12/2015 |
| (3)<br>Blue<br>inhibition | 9 | m-c | Y2 only & Y2 mixed with<br>+V2, +V3, +B1, +B2, +C | 50 + 5 | 10 | 1 | 10 | 150 | 08/05/2015 |
|  | 10 | m-c | Y2 mixed with<br>+V2, +V3, +B1, +B2, +C | 50 + 2.5 | 21 | 3 | 7 | 150 | 09-12/05/2015 |
|  | 11 | d-c | +B2 vs.<br>+V2, +V3, +B1, +C | 50 + 2.5 | 10 | 5 | 8 | 150 | 19-23/05/2015 |
| (4)<br>UV<br>response | 12 | m-c | UV(1, 2, 3, 4), V(1, 2) | 40 | 20 | 4 | 5 | 150 | 22-28/09/2015 |
|  | 13 | m-c | UV(1, 3, 4), V(2, 3), B1 | 40 | 20 | 4 | 5 | 150 | 06-10/10/2015 |
|  | 14 | d-c | UV1 vs.<br>UV3, UV4, V2, V3 | 40 | 10 | 10 | 4 | 150 | 12/10-02/11/2015 |

\* m-c = Multiple-choice experiment, d-c = Dual-choice experiment / \*\* LED colour abbreviations according to Table 1 / \*\*\* Trial duration varied by $\pm 0.10$ h at most

Wavelength-dependent responses were initially investigated in multiple-choice experiments and subsequently relevant LEDs were selected and tested in dual-choice experiments to determine standardized spectral efficiencies. All multiple-choice and dual-choice experiments were performed in 2015. The experiments took place in the described choice arena and replicates were conducted in consecutive trials on different daytimes and days. Trials were conducted between 10:00 and 17:00 h. Experiments regarding the ‘settling response’ were conducted from February to May. With increasing day length and brighter ambient light conditions in the greenhouse, whiteflies orientated more readily to the traps, hence trial durations could be reduced and number of trials per day could be increased within this time. UV response experiments were conducted from September to November but suffered from weaker responses and low recapture rates and trial durations were adjusted accordingly (Table 2).

In multiple-choice experiments, the LED trap screens in question (six or five) were placed in the holes of the choice arena background and the order was changed randomly for each replication. 150 or 200 whiteflies were released per replication and the number of trapped individuals on each trap were counted after a given period (0:30 – 1:30 h). Afterwards the cage was cleaned carefully from remaining whiteflies with a handheld vacuum cleaner before starting the next trial. The procedure for dual-choice experiments was similar, but four holes for trap screens were covered with black plastic film and only the two inner holes were equipped with the two LED traps. Again, trap positions were changed randomly for each replication. The measurement of ambient light conditions were averaged over each experimental trial and considered in the dataset as co-variable.

### Block 1: Green response experiments (Exp. 1-5)

The wavelength dependence of the ‘settling response’ in the green-yellow range was studied using 12 LED colours at equal photon fluxes including the adjacent blue and red ranges. Four multiple-choice experiments were conducted comparing six LEDs simultaneously in one experiment. Experiment 1 compared LEDs ranging from blue to green and exp. 2 those ranging from green to red. These experiments were interlinked by one green LED (G3) presented in both experiments. Then the most targeted LEDs from these two experiments were selected and compared in exp. 3. Here, the sex ratio of the trapped whiteflies on each LED colour was also determined in five of the 20 replicates (last trial of each day). Finally, the previously less preferred blue and red LEDs were compared separately in exp. 4.

The most attractive chartreuse green LED (G4 - 550 nm centroid wavelength) from the multiple-choice experiments was selected as a reference to determine standardized spectral efficiencies of seven LED colours (test lights) from blue-green to amber (BG, G1-3, Y1-2, A) and successively tested against the green reference LED in dual-choice assays (exp. 5). The responses were the relative choice frequencies on the test lights which were graphically displayed relative to the reference light which was set to maximum response. The spectral efficiencies of the tested LED colours were normalized to obtain a standardized LED based action spectrum of the ‘settling response’ under daylight conditions. The experiment was conducted with one replicate per colour per day and a randomized order of the colours per day.

### Block 2: Intensity dependences (Exp. 6-8)

Following the determination of the spectral efficiency in the ‘settling response’ (see exp. 5) the intensity dependence of the choice behaviour was determined in the same dual-choice setup (exp. 6). The intensity of the chartreuse green reference light (G4) was reduced by 50% and tested against four spectrally adjacent green and yellow LEDs (G1, G3, Y1, Y2). The data of this experiment were merged with the initial data of these colours (exp. 5, LEDs at equal intensity) to illustrate the intensity-dependent changes in the spectral efficiencies.

The influence of different intensities of the same colour on the preference in a multiple-choice setup was looked at in another experiment with six yellow (Y2) LED traps at different intensities (exp. 7). One trap was set to maximum intensity and intensities of the others were reduced evenly.

In a final multiple-choice experiment, the same yellow LED traps were tested at equal intensities with randomized order to evaluate the bias regarding their positions in the choice arena (exp. 8).

### Block 3: Blue inhibition experiments (Exp. 9-11)

A potential inhibitory blue-green chromatic mechanism was studied combining five panels with yellow LEDs (Y2 - 590 nm centroid wavelength) with two violet LEDs (V2 - 415, V3 - 435 nm), two blue LEDs (B1 - 447, B2 - 469 nm), and one cyan LED (C - 500 nm), respectively. Yellow LEDs were used here because we assume that they stimulate mostly the green receptor on the long wavelength side to ensure that inhibitory interaction effects can be attributed to the mixture with blueish LEDs. One additional panel remained with only yellow LEDs and the intensity of all six yellow LED panels was set to 50 µmol m^-2^ s^-1^ on the trap screen. A small amount of 5 µmol m^-2^ s^-1^ (= 9.1% relative intensity) of the respective blueish LED light was added.

In a first multiple-choice experiment, the five LED trap screens with yellow-blueish mixture and the pure yellow LED trap were compared (exp. 9). The pure yellow LED trap consequently had a 9.1% lower total intensity due to the lack of additional blueish light. In a second multiple-choice experiment (exp. 10), the pure yellow LED trap was excluded from the setup and the intensities of blueish LEDs were further reduced to 2.5 µmol m^-2^ s^-1^ (= 4.8% relative intensity).

The most unattractive yellow-blue combination (Y2+B2) was selected as reference to determine standardized spectral efficiencies of the other four yellow-blue combinations (test lights) in successive dual-choice assays (exp. 11). Here, the responses were the relative choice frequencies on the reference light, representing a measure of inhibition. A standardized LED based action spectrum of ‘settling inhibition’ was constructed according to the procedure in the green response experiments. The experiment was conducted with two replicates per colour per day and randomized order of the colours within the day.

### Block 4: UV response experiments (Exp. 12-14)

The wavelength dependence of the ‘migratory behaviour’ in the UV range was studied using eight LEDs from UV to blue at equal photon fluxes. The first multiple-choice experiment compared LEDs from the narrow UV to violet range (exp. 12). In the second multiple-choice experiment, the spectral range was extended to blue with larger spectral steps between the LED colours (exp. 13).

The most attractive UV LED (UV1 - 373 nm centroid wavelength) was selected as reference to determine the standardized spectral efficiencies of four LED colours (test lights) from UV to violet (UV3, UV4, V2, V3) in dual-choice assays (exp. 14). A standardized LED based action spectrum of the UV response was constructed according to the procedure in the green response experiments (see Block 1). The experiment was conducted with two replicates per colour per day and randomized order of the colours within the day.

### Colour choice models

An empirical colour choice model was built to describe the wavelength preference in the ‘settling response’ based on opponent chromatic interaction of a green and a yet undescribed blue photoreceptor. Modelling of the UV response was performed assuming achromatic processing based only on the UV receptor. As no reliable data of photoreceptor sensitivities are available for whiteflies, the peak sensitivities were approximated by this method.

Photoreceptor sensitivity templates (Govardovskii et al., 2000) were fitted for different photoreceptor peak sensitivities of a putative UV, blue, and green photoreceptor, respectively. The peak sensitivities of the green and the blue receptor were altered in 5 nm steps in the range of 500 - 545 nm (green) and 470 - 495 nm (blue) resulting in 60 potential combinations. The peak sensitivities of the UV receptor was changed in 10 nm steps in the range of 340 - 370 nm.

The photon catch *P* of a photoreceptor can be calculated with the photoreceptor sensitivity function *S*(λ) and the spectrum of the (LED) stimulus light *I*(λ) (Kelber et al., 2003):
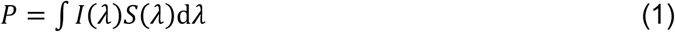

The photon catches of each LED colour (and its combinations) were calculated for each potential photoreceptor position. Photoreceptor excitations *E* were calculated from photon catch values using a nonlinear transformation (Chittka, 1996):
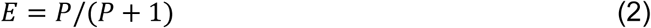

This resulted in excitation values for each LED and each photoreceptor (*E*_UV_, *E*_B_, *E*_G_) at varying positions. The excitations of the colour opponent mechanism *E*_opp_ were calculated as difference between green and blue photoreceptor excitations:
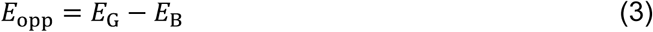

These values were connected to the LED choice datasets of the ‘green response’, the ‘blue inhibition’ and the ‘UV response’, resulting in three separate models.

For the ‘green response model’ the mean relative choice frequencies from the ‘green response experiments’ (exp. 1-3, 5) were combined and plotted against *E*_opp_ values of each receptor configuration. The data from multiple-choice experiments were thereby normalized to the most attractive chartreuse green LED (G4). The first dataset was built based on the outcome of exp. 1 and 2 which were connected via the linking green LED (G3) used in both experiments. Exp. 3 was taken as second dataset and the normalized spectral efficiencies from exp. 5 as third dataset. A preference restriction was implemented which considers that the highest excitation value should correspond with the most attractive chartreuse green LED (G4):
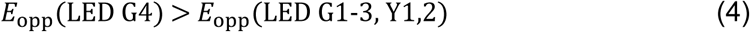

For the ‘blue inhibition model’ the data from the ‘blue response experiments’ with mixed yellow and blue LEDs (exp. 10, 11) were plotted against *E*_opp_ values. Here, the indirect response was the inhibition of the attraction and the highest response was referred to the most inhibiting blue LED. Therefore, the mean relative choice frequencies from the multiple-choice experiment (exp. 10) were inverted and normalized to the most unattractive yellow-blue combination (Y2+B2). The normalized spectral efficiencies of inhibition from exp. 11 were taken as second dataset. Here, the lowest excitation value should correspond with the blue LED (B2) inhibiting the attraction towards yellow LEDs the most:
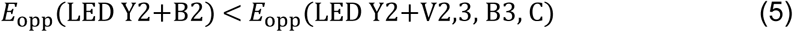

For the ‘UV response model’, achromatic processing based solely on the UV receptor was assumed. Therefore, the excitation values *E*_UV_ were directly plotted against the normalized relative response data from the multiple-choice experiments (exp. 12, 13) and the dual-choice spectral efficiency experiment (exp. 14). The restriction that the highest excitation value should correspond with the most attractive UV LED is described by:
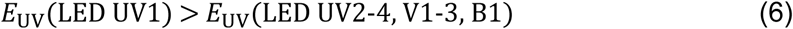

All models’ significant linear regressions (α = 0.05) fulfilling the preference restrictions were fitted and the models were assessed based on R^2^ values. All analyses and graphical display related to the colour choice models were performed in Microsoft Excel 2016.

### Statistical analysis

The statistical analyses were performed in R (Version 3.2.1; R Core Team, 2015).

The multiple-choice experiments (exp. 1-4, 7-8, 9-10, 12-13) were analysed with linear models using the lm() function. The response variables were the ln(x + 1) transformed numbers of trapped whiteflies on each LED trap. In colour choice experiments, the explanatory variable was the LED colour. The ambient light intensity (visible light or UV radiation) measured throughout the experiments was included as co-variable for the experiments of the green and UV response. Initial Block factors (day, daytime) of the consecutive experiments were excluded after model selection using Akaike’s Information Criterion (Burnham and Anderson, 2010). Interactions between the colour and the ambient light intensity were included in the analyses of the green response experiments 2 and 3. Separate linear models were fitted to analyse the total numbers of trapped whiteflies in the given time dependent on the ambient light intensity. In the analyses of the multiple-choice experiments with different LED intensities (exp. 7), LED trap intensity and its interaction with ambient light intensity were explanatory variables. In the analyses of the multiple-choice experiment with equal LED intensities (exp. 8), the individual LED trap number and the position in the choice arena were the explanatory variables. ANOVA was used to determine influences of explanatory variables and interactions in the linear models. Tukey-type pairwise comparisons regarding LED colours and intensities were performed at α=0.05 using the lsmeans package (Lenth, 2015).

The sex ratio in the multiple-choice experiment 3 was analysed with a generalized linear model using the glm() function with binomial distribution and logit link. The response variable was the odds ratio between males and females on each trap and the explanatory variable was the colour. The dual choice experiments (exp. 5, 6, 11, 14) were analysed with generalized linear models (quasibinomial, logit link). The response variable was the odds ratio between the number of trapped individuals on test and reference LED traps. Explanatory variable was the respective colour of the test LED. The ambient light intensity was included as co-variable for the spectral efficiency experiments on green and UV response. An interaction between colour and ambient light was further included in the green response analysis. Deviance analyses were performed to determine influences of explanatory variables and interactions in the generalized linear models. In the intensity dependence dual choice experiment (exp. 6), pairwise comparisons were performed between intensity levels (α=0.05, lsmeans package). User-defined interaction contrasts were created to compare intensity-dependent changes of choice frequencies between colours using the package statint (Kitsche and Schaarschmidt, 2015). Tukey-type comparisons on interaction contrasts were performed using the multcomp package (Hothorn et al., 2008). Graphs were created using the ggplot2 and gridExtra package (Wickham, 2016; Auguie, 2012).

## Results

### Block 1: Green response experiments (Exp. 1-5)

#### Experiment 1

The results showed hardly any response of whiteflies to the blue (B2 - 469 nm), cyan (C - 500 nm), and blue-green (BG - 512 nm) LED, and a steep significant increase in the preference among green LEDs (G1-3) with only slightly different centroid wavelengths of 524, 528, and 533 nm (Fig. 3A).

**Fig. 3.**
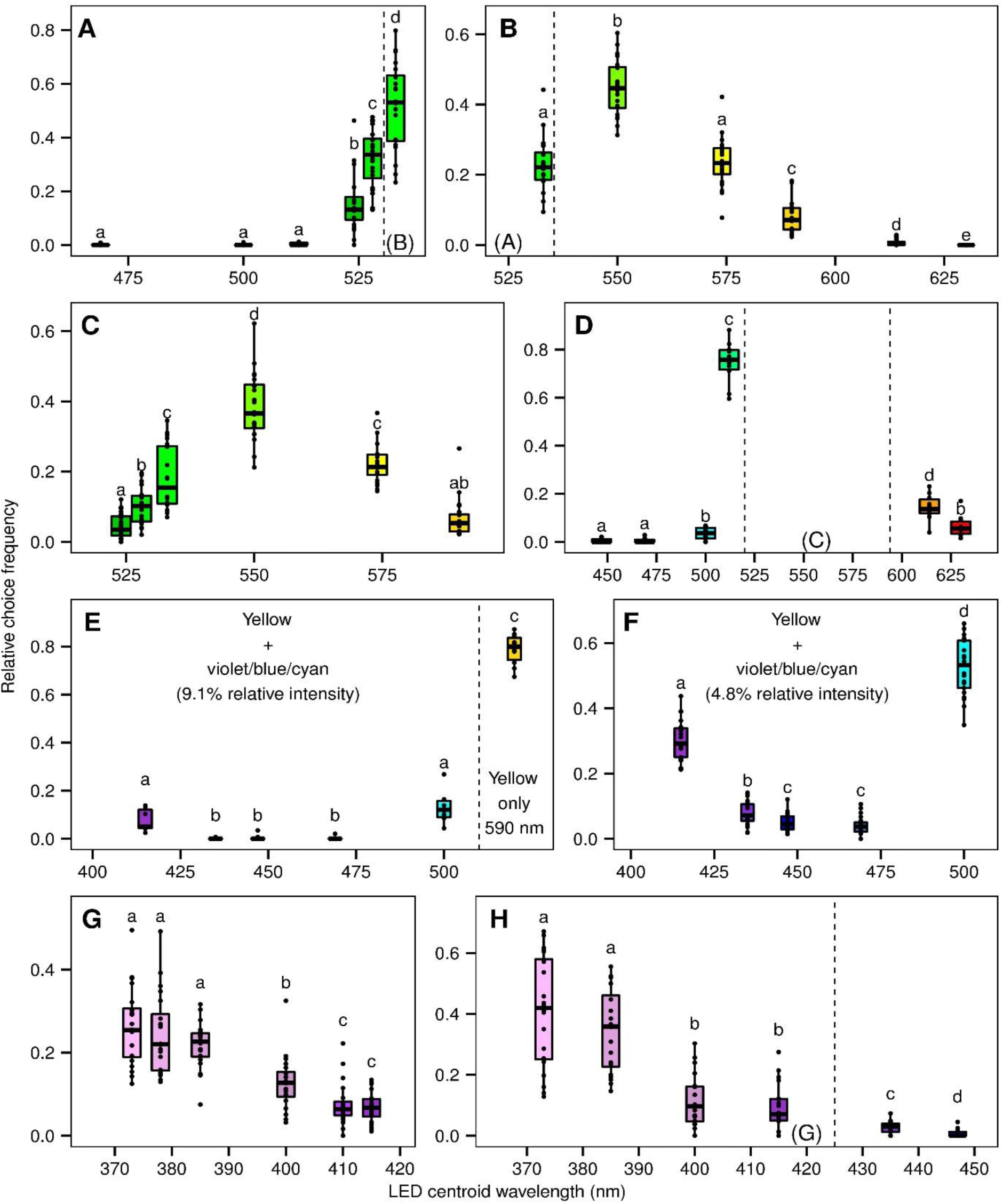
Wavelength preferences of *Trialeurodes vaporariorum* in LED multiple-choice experiments. (A) blue - green (exp. 1), (B) green - red (exp. 2), (C) green - yellow (exp. 3), (D) blue & red (exp. 4), (E) yellow + violet - cyan & pure yellow (exp. 9), (F) yellow + violet - cyan (exp. 10), (G) UV - violet (exp. 12), (H) UV - blue (exp. 13). See Table 2 for experimental details. Dashed vertical lines and panel letters in brackets on the bottom indicate spectral overlapping of the experiments. Dots show original data points. Boxes indicate interquartile ranges (IQR) with median (thick line). Whiskers comprise values within 1.5 × IQR. Different letters indicate significant differences of the factor LED colour within each experiment (Linear Model: ln (x+1) ~ LED colour × ambient light intensity, Tukey post-hoc tests, P=0.05).

No significant influence of the ambient light or the interaction with colours were observed in the fitted linear model (Fig. 4A). This indicates that whiteflies discriminated green LEDs over the whole ambient light intensity range. The overall recapture rate was 69.0 ± 6.6% (Mean ± s.d.) within 1:15 ± 0:10 h. A separately fitted linear model shows no significant increase of the total recaptures with rising ambient light intensity.

**Fig. 4.**
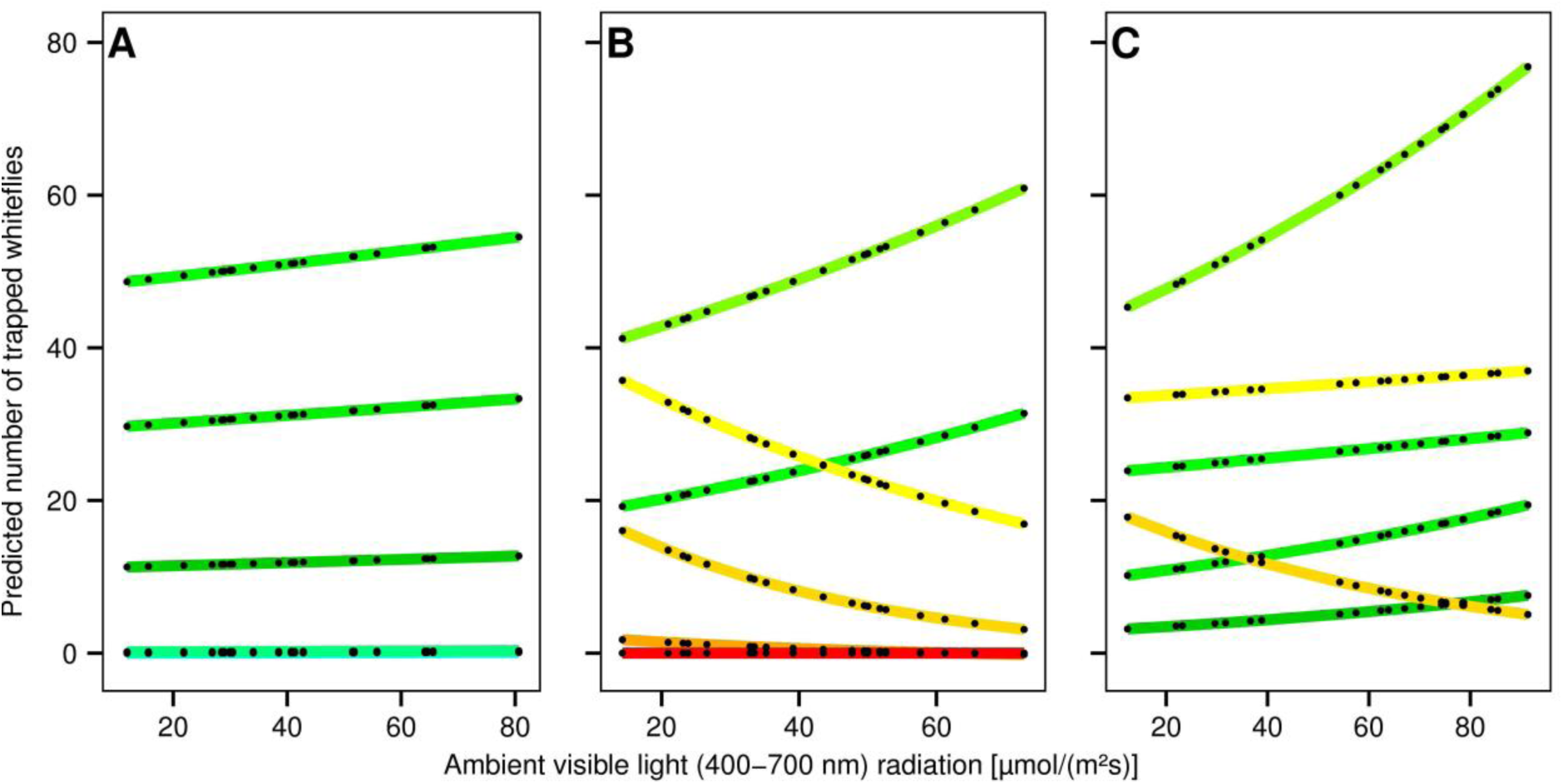
Interactions of wavelength preferences in *Trialeurodes vaporariorum* with ambient light intensity in LED multiple-choice experiments based on the fitted models. Black dots connected by coloured lines show backtransformed predictions from the linear models (ln(x+1) ~ LED colour × ambient light intensity) fitted to the data of (A) exp. 1, (B) exp. 2, (C) exp. 3. See Fig. 3A, B, C for original data and associated line colours (lines at zero partly overlay each other).

#### Experiment 2

In the green range, the preference further increased revealing chartreuse green (G4) with 550 nm centroid wavelength as the most attractive LED (Fig. 3B). Towards the yellow spectrum with the two yellow LEDs (Y1 - 574, Y2 - 590 nm), the preference declined and only a weak response to amber (A - 614 nm) and no response to the red (R - 630 nm) LED were noticed.

In contrast to exp. 1, a significant influence of ambient light and the interaction with colour were observed (both P<0.001). At darker conditions, the response to yellow was relatively stronger while the corresponding response to green was weaker (Fig. 4B). With increasing ambient light intensity, the response to green LEDs (G3, G4) increased while the response to yellow LEDs (Y1, Y2) decreased correspondingly. This resulted in a cross-over interaction between the second most attractive green (G3) and yellow (Y1) LEDs which are on average of similar attractiveness but with increasing ambient light intensity G3 became more attractive. The overall recapture rate was 75.6 ± 10.9% (mean ± s.d.) within 1:15 ± 0:10 h and total recaptures were not influenced by ambient light.

#### Experiment 3

When the selected attractive green and yellow LEDs (G1-4, Y1-2) were compared, the results show that the relative preferences resemble an action spectrum (Fig. 3C).

No significant influence of the ambient light but a significant interaction with the colour could be determined (P=0.019). At darker conditions, the preferences were more evenly distributed across all colours and with rising ambient light intensity the preference was pointed more towards the most attractive chartreuse green LED (G4) while the preference towards the second most attractive yellow (Y2) decreased (Fig. 4C). The overall recapture rate was 82.8 ± 10.0% (mean ± s.d.) within 1:15 ± 0:10 h. The totally recaptured numbers increased significantly with rising ambient light (P=0.003), primarily due to the strongly increasing preference for the most attractive chartreuse green (G4).

The ratio of females on the LED colours were 68% on G1, 72% on G2, 72% on G3, 72% on G4, 81% on Y1, and 81% on Y2; the overall ratio was 74.5%. The ratio of females was slightly higher on the yellow LEDs but statistically no significant effect of LED colours on the sex ratio was observed (GLM, Analysis of Deviance, P=0.16).

#### Experiment 4

When the previously attractive range was excluded, whiteflies significantly preferred the blue-green (BG - 512 nm) LED and only few landings were recorded on cyan (C - 500 nm), amber, and red LED traps (Fig. 3D). The overall recapture rate was 41.9 ± 10.6% (mean ± s.d.) within 1:15 ± 0:10 h.

#### Experiment 5

In the spectral efficiency dual-choice experiment the response declined steeply over the three green LEDs to very little relative response towards the blue-green LED. On the long wavelength side, the response declined a bit wider over the two yellow LEDs to almost zero response on the amber LED. The obtained action spectrum was similar to the action spectra derived from previous multiple-choice experiments (Fig. 5). No significant influence of the ambient light but a significant interaction with colour could be determined (GLM, Analysis of Deviance, P=0.005). The recapture rate was 82.0 ± 13.5% (mean ± s.d.) within 0:40 ± 0:10 h.

**Fig. 5.**
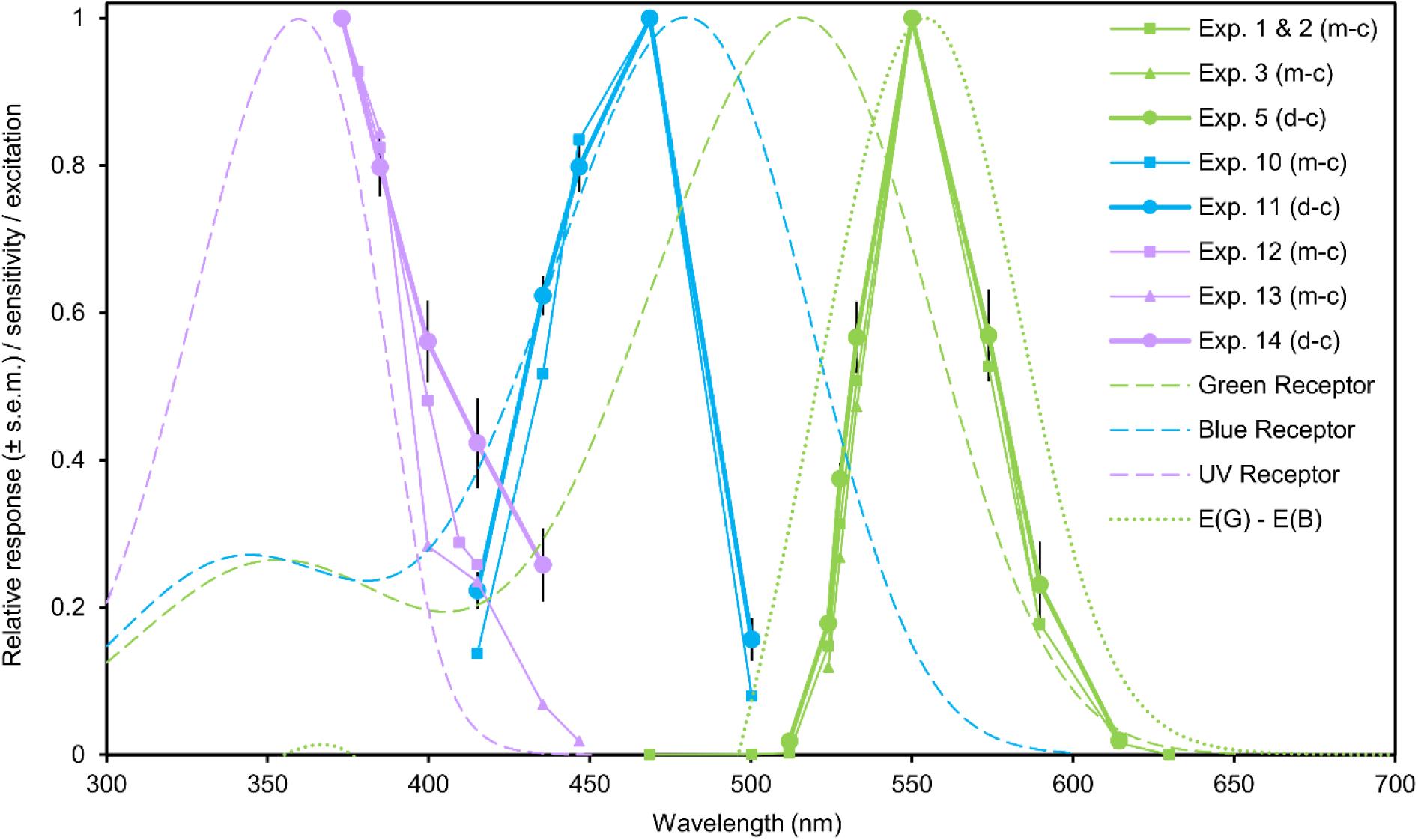
Spectral efficiencies of *Trialeurodes vaporariorum* derived from LED choice experiments and modelled putative photoreceptor sensitivities with resulting theoretical action spectrum of the ‘settling response’ based on blue-green opponency. Coloured symbols connected by solid lines show relative responses in the respective spectral range dependent on LED centroid wavelengths (see Table 1 and Fig. 1 for LED centroid wavelengths and spectra). Green data points refer to the green response (‘settling’) and violet data points indicate the UV response. Blue data points are derived from mixing experiments with equal yellow and different blueish LEDs indicating the ‘settling inhibition’ as an inverse response to blue. Thin solid lines with squares or triangles show normalized mean relative choice frequencies from multiple-choice (m-c) experiments. Thick solid lines with circles (± s.e.m) show normalized mean relative choice frequencies from dual-choice (d-c) spectral efficiency experiments. See Table 2 for experimental overview and Fig. 3 for original data in multiple-choice experiments. Dashed coloured lines show photoreceptor sensitivity templates (Govardovskii et al., 2000) with peak sensitivities at 360 nm (UV), 480 nm (blue), and 515 nm (green) estimated in colour choice models. The dotted green line describes the modelled blue-green opponency as difference of photoreceptor excitations and represents the theoretical action spectrum of the ‘settling response’.

### Block 2: Intensity dependencies (Exp. 6-8)

#### Experiment 6

When the intensity of the green reference light was reduced following the determination of the spectral efficiency (exp. 5), the choice frequencies on the respective green and yellow LEDs increased significantly (G1, G3, Y1: P<0.001; Y2: P=0.003; Fig. 6A). The increase was strongest on G1, thereby almost reaching equal response (choice frequency=0.5, Logit=0, indicated as dashed line in Fig. 6A) as on the reference LED (G4). The strength of increase was slightly lower on G3 and Y1 but the choice frequencies reached an even higher level than on the reference LED. The increase in attractiveness was significantly lower on Y2 compared to the other LEDs (Y2 vs. G1, G3: P<0.001; Y1 vs. Y2: P=0.015), and the response remained below the corresponding response to the reference LED.

**Fig. 6.**
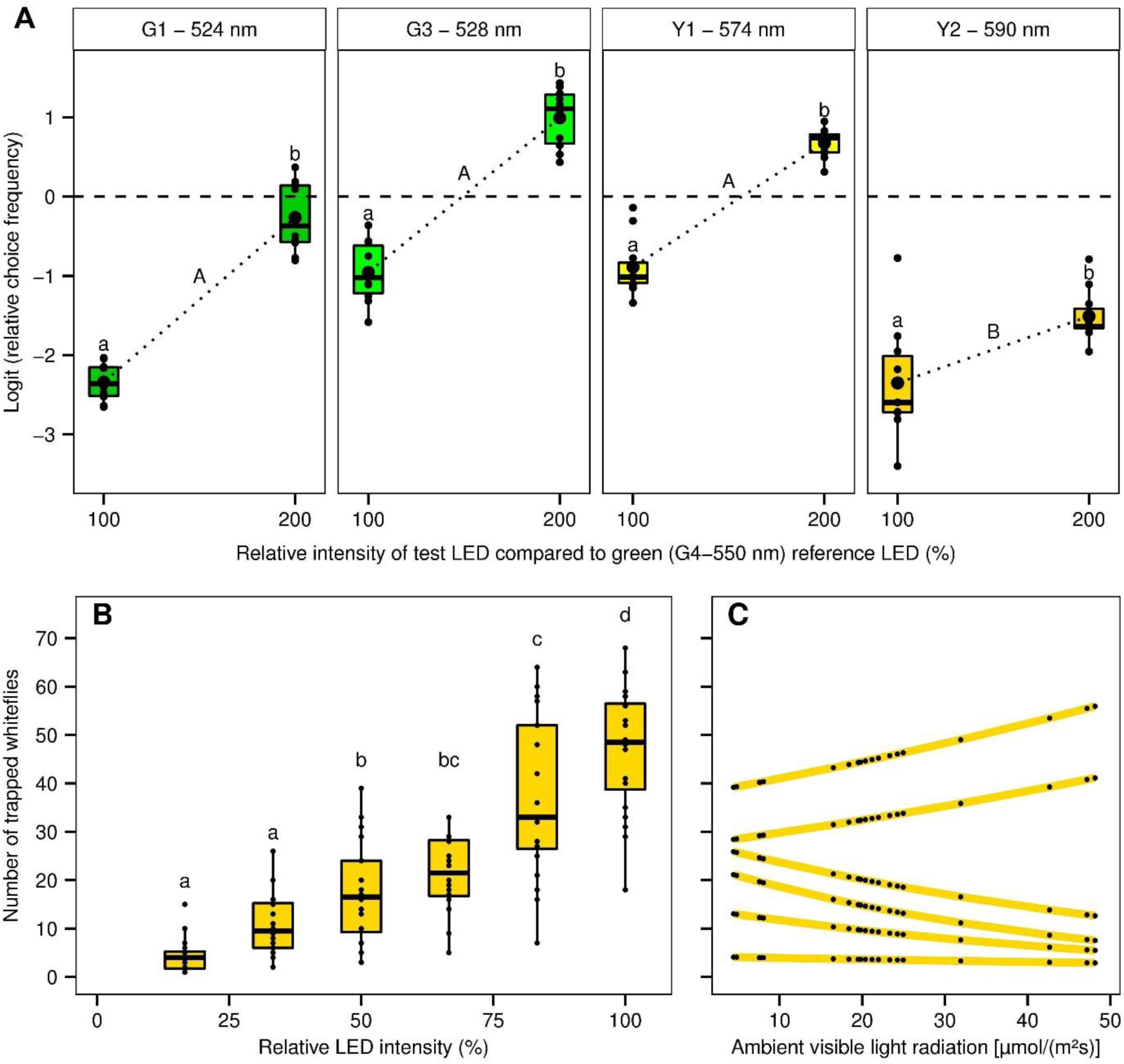
Intensity dependencies in the colour choice behaviour of *Trialeurodes vaporariorum*. (A) Intensity dependences between spectral efficiencies. Panels show Logit-transformed choice frequencies in dual-choice experiments with four LED colours at equal (=100%) and double (=200%) relative intensity of the reference LED (G4 - 550 nm). Relative intensity changes were created by reducing the reference LED intensity by 50%. The dashed horizontal line indicates equal choice frequencies (=0.5, Logit=0) of test and reference LED. The dotted line connects mean choice frequencies of intensity levels. Different small letters indicate significant differences between intensity levels within each colour (GLM, pairwise comparisons, P=0.05). Capital letters indicate significant differences of intensity-dependent changes of choice frequencies between colours (GLM, user defined interaction contrasts, P=0.05). (B) Intensity Preference for equal yellow LEDs with 590 nm centroid wavelength at different relative intensities. Different letters indicate significant differences of the factor LED intensity (Linear Model, Tukey post hoc test, P=0.05). (C) Corresponding interaction with ambient light intensity based on the fitted model. Black dots connected by yellow line show predicted values from the linear model (Linear Model: ln (x+1) ~ LED intensity × ambient light intensity).

#### Experiment 7

Different intensities of the same yellow (Y2) in a multiple-choice experiment showed the strongest response on the brightest LED and a constant decrease of attractiveness towards the lowest intensity (Fig. 6B).

A significant influence of ambient light intensity on the trapped numbers on each colour was observed (P=0.048, Fig. 6C). The interaction between ambient light and LED intensity was an explanatory factor according to model selection by AICs (P=0.079).

#### Experiment 8

When the yellow LEDs from the previous experiment were compared at equal intensities, the LED position had a significant influence on the numbers trapped (P=0.018, data not shown). More whiteflies were trapped on the outer side positions compared to the inner positions. But due to randomization and repetitions this effect could be neutralised resulting in no significant effect on the trapped numbers on respective LED traps (P=0.28).

### Block 3: Blue inhibition experiments (Exp. 9-11)

#### Experiment 9

Most of the whiteflies were trapped on the LED trap with pure yellow (Y2 - 590 nm). Little response was obtained when yellow was additively combined with small intensities of the shortest wavelength violet (V2 - 415 nm) or the longest wavelength cyan (C - 500 nm) LED. Almost no trappings were recorded on the combinations with the intermediate violet (V3 - 435 nm) and blue (B1 - 447, B2 - 469 nm) LEDs. The results clearly indicate that the “settling response” was inhibited by blueish light (Fig. 3E). The overall recapture rate was 92.8 ± 4.9% (mean ± s.d.) within 0:30 ± 0:10 h.

#### Experiment 10

When the pure yellow light was excluded from the setup and the intensity of blueish light was further reduced, the preferences exhibited in the previous experiment were clearly emphasized. Highest trap catches were recorded on the yellow-cyan combination and lowest catches on the yellow-blue combinations (B1 - 447, B2 - 469 nm). The preference increased again for the adjacent violet (V3 - 435 nm) and for the shortest wavelength violet (V2 - 415 nm) LED in particular. The data resemble an inverse action spectrum of inhibition of the ‘settling response’ (Fig. 3F). The overall recapture rate was 89.7 ± 10.5% (mean ± s.d.) within 0:30 ± 0:10 h.

#### Experiment 11

On the short wavelength side, the inhibition declined successively from UV to blue (B1) and violet (V2, V3) LEDs. On the long wavelength side, the inhibition strongly decreased in one big step to the cyan (C) LED. Again, the obtained action spectrum was quite congruent with the one derived from the multiple-choice approach (Fig. 5). The recapture rate was 75.4 ± 13.0% (mean ± s.d.) within 0:30 ± 0:10 h.

### Block 4: UV response experiments (Exp. 12-14)

#### Experiment 12

The highest responses were recorded on the first three UV-A LEDs (UV 1-3) with closely related centroid wavelengths of 373, 378, and 385 nm but these preferences did not differ among each other. The preference declined over 400 nm (UV4) to the violet (V1 - 410, V2 - 415 nm) LEDs which showed the lowest but still detectable response (Fig. 3G).

A significant influence of the ambient UV radiation on the trapped numbers on the colours was observed in the fitted linear model (*p*=0.003). The overall recapture rate was 46.8 ± 10.7% (mean ± s.d.) within 1:30 ± 0:10 h. A separately fitted linear model showed that the totally recaptured numbers decreased with rising UV radiation intensities (P=0.006).

#### Experiment 13

When the tested spectral range was extended to blue, the preference further declined on the long wavelength violet (V3 - 435 nm) and very low responses were still detected on the short wavelength blue (B1 - 447 nm) LED (Fig. 3H).

UV radiation had a significant influence on the trapped numbers on the colours (P=0.046). The overall recapture rate was 46.8 ± 10.7% (mean ± s.d.) within 1:30 ± 0:10 h and total numbers were not significantly influenced by ambient UV radiation.

#### Experiment 14

The response declined successively over the tested UV and violet colours but was still quite prominent on the long wavelength violet (V3). The obtained half-sided action spectrum was wider and not entirely congruent with the ones derived from the previous multiple-choice experiments (Fig. 5). The recapture rate was 23.6 ± 10.0% (mean ± s.d.) within 1:30 ± 0:10 h.

### Colour choice models

In the ‘green response model’ and the ‘blue inhibition model’, several combinations of blue and green photoreceptor peak combinations led to significant linear regressions which fulfil the preference restrictions (Table 3).

**Table 3.**
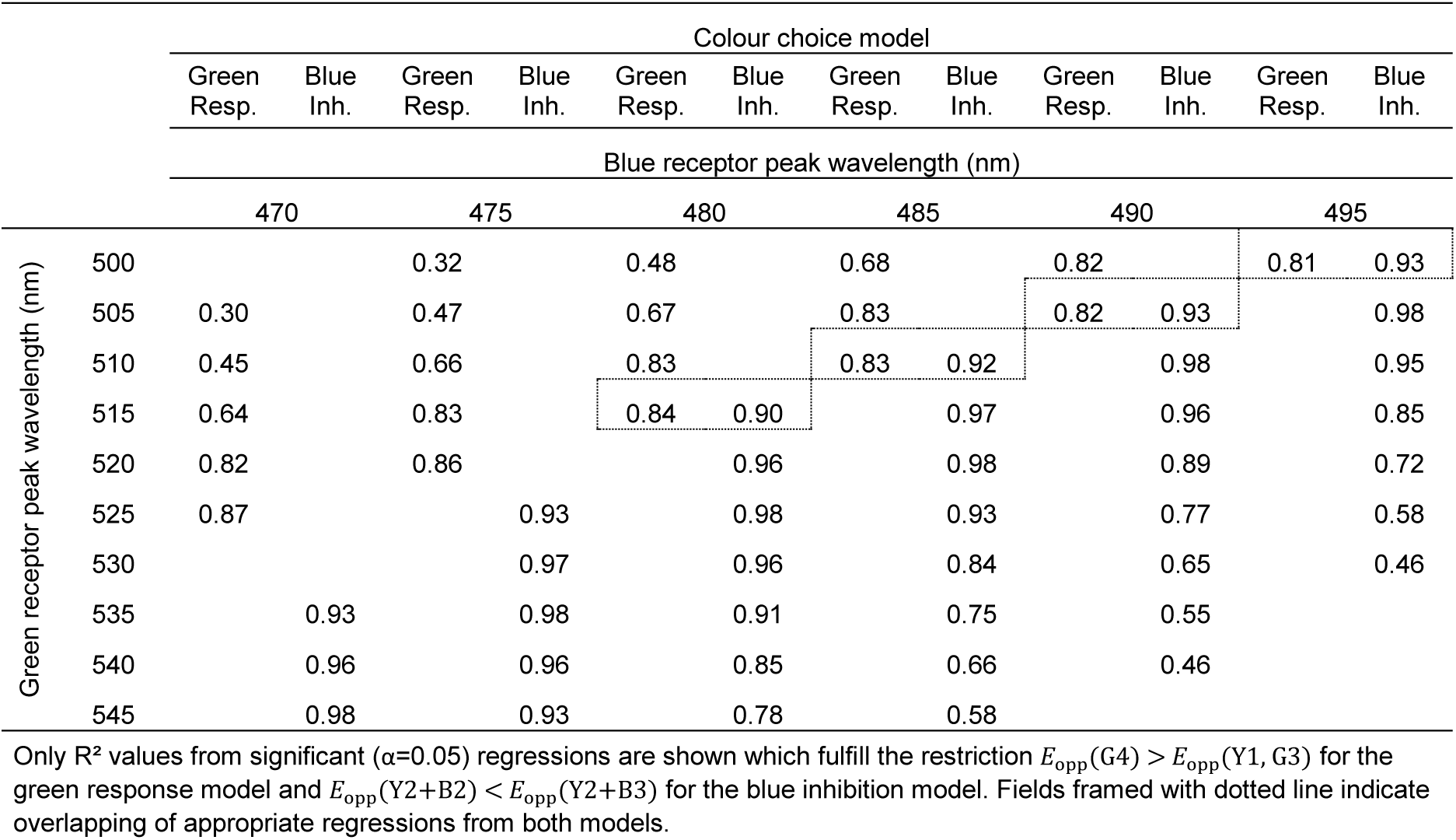
R^2^ values from linear regressions of colour choice models for different photoreceptor combinations based on blue-green opponency.

For the ‘green response model’, regressions with good fits (R^2^ ≥ 0.8) were found for receptor peak combinations from 470 & 525 nm at widest distance to 495 & 500 nm at lowest distance from each other. In the blue inhibition model, good fits (R^2^ ≥ 0.9) were found for combinations from 470 & 545 nm at widest distance to 495 & 500 nm at lowest distance. Well-fitting regressions which fulfil the restrictions in both models overlap at receptor combinations of 480 & 515 nm, 485 & 510 nm, 490 & 505 and 495 & 500 nm (Tab. 3). Selected regressions for potential blue and green receptor peaks at 480 and 515 nm are shown in Fig. 7A,B. The modelled potential photoreceptors based on template formulas (Govardovskii et al., 2000) and the resulting theoretical relative action spectrum of the ‘settling response’ based on blue-green opponency are shown in Fig. 5.

**Fig. 7.**
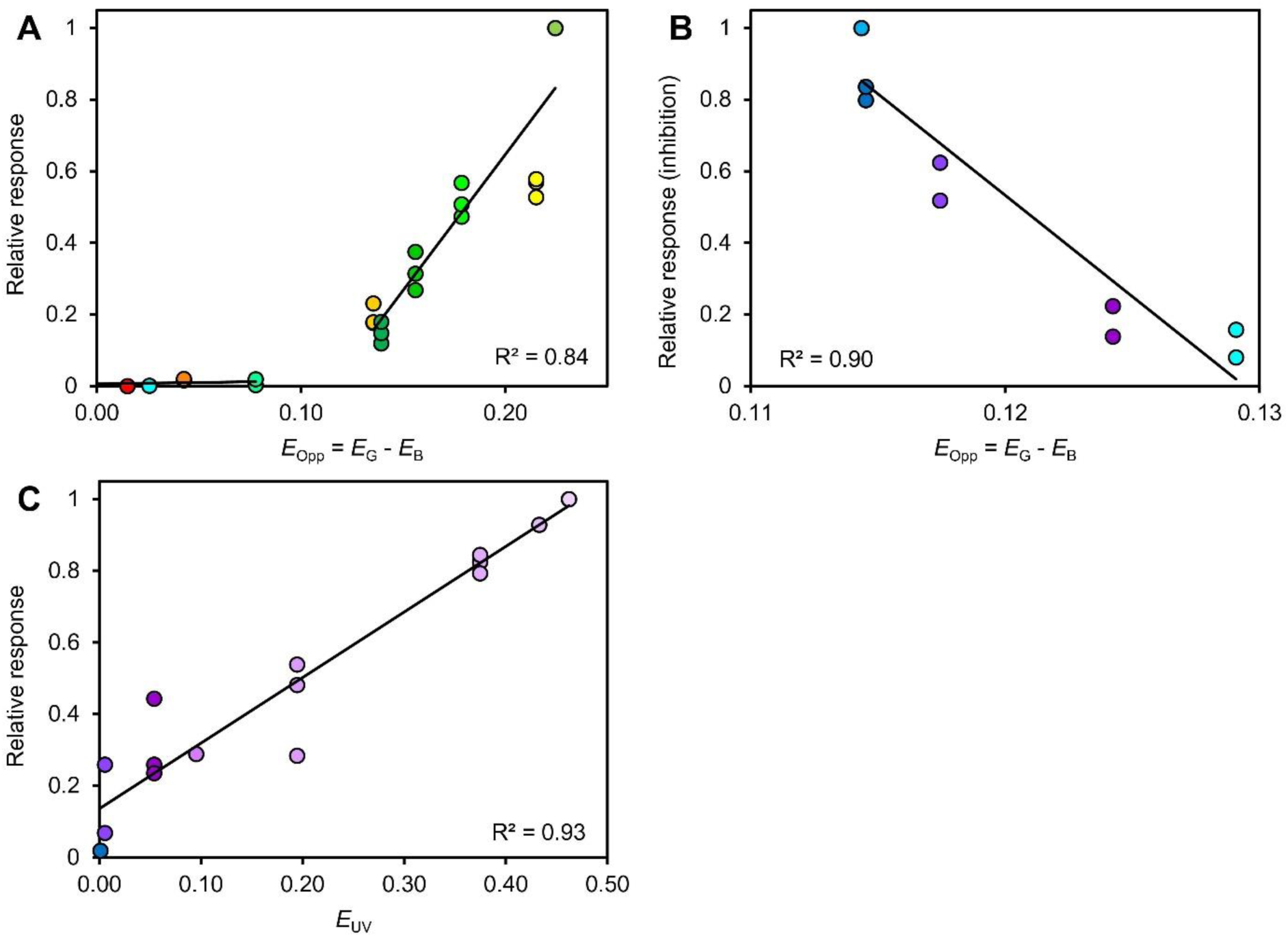
Selected linear regressions of colour choice models. (A) Green response model and (B) Blue inhibition model for photoreceptor peaks at 480 nm and 515 nm based on blue-green opponency (See Table 2) (C) UV response model with photoreceptor peak at 360 nm.

Fig. 7C shows the best fitting linear regression of the ‘UV response model’ with a photoreceptor peak at 360 nm (R^2^ = 0.93) and the modelled receptor is also shown in Fig. 5. The restriction that the highest excitation value corresponds with the most attractive UV LED is also fulfilled for adjacent receptor peaks at 340, 350, and 370 with R^2^ values of 0.77, 0.90, and 0.86, respectively.

## Discussion

### Main findings

This study reveals that *Trialeurodes vaporariorum* possesses a yet undescribed photoreceptor sensitive towards blue light and an inhibitory blue-green chromatic mechanism which controls a ‘wavelength-specific behaviour’ referred to as ‘settling response’ (Coombe, 1981). Besides this chromatic processing, the behavioural control is distinctly intensity-dependent. The known response to UV radiation based on a UV sensitive photoreceptor related to migratory behaviour could also be confirmed in our study (Coombe, 1981; 1982). As a consequence, we could conclude that *T. vaporariorum* possesses a trichromatic visual system.

### Wavelength dependence of the ‘settling response’ and interaction with ambient light

The chartreuse green LED with 550 nm centroid wavelength proved to be most attractive (Fig. 3B,C) and consequently constitutes the peak of the LED based action spectrum of the ‘settling response’ (Fig. 5). This meets our expectations as it is in line with earlier studies from MacDowall (1972) and Coombe (1981) also showing action spectra peaking at 550 nm. As only this LED was available in the region between 533 and 574 nm, it is possible that the actual peak slightly differs which is also possible for both reported studies which used monochromatic light in wide steps of 10 and 50 nm. When only one receptor controls the behaviour, the action spectrum should roughly exhibit the shape of the underlying receptor (Skorupski and Chittka, 2011). But our action spectrum as well as the reported data are more narrowly tuned to the green-yellow range and shifted to the longer wavelength range compared to the spectral efficiency peak at 520 nm which was determined by ERG recordings by Mellor et al. (1997). This discrepancy suggests the involvement of opponent processing and the extraction of chromatic signals (Skorupski and Chittka, 2011). Nevertheless, from an evolutionary perspective it seems natural that these action spectra peak around 550 nm which corresponds quite accurately with the peak reflectance and transmittance of green leaves, corroborating the fact that the visual systems of herbivores are adapted to host plant detection (MacDowall, 1972; Döring et al., 2009; Prokopy and Owens, 1983; Kelber and Osorio, 2010).

An important observation with regard to potential chromatic processing was that green LEDs with similar spectra of only 4-5 nm difference could be differentiated by *T. vaporariorum* as shown by the multiple-choice experiments (Fig. 3A,C). Moreover, the discrimination was exhibited consistently over the whole range of ambient light intensity, whereas yellow LEDs were to some extent confused with green ones at darker conditions (Fig. 4). Compared to naturally reflecting objects, the constant intensity of LED light is uncoupled from illuminating light intensity and should theoretically appear as brighter or darker in relation to changing ambient light intensity. Colour vision is defined as the ability to detect spectral variations in the light independent of the intensity (Kelber et al., 2003). Photoreceptors adapt to the intensity of perceived light versus the background light by adjusting their responses through various mechanisms (Laughlin and Hardie, 1978; Arshavsky, 2003; Warrant and Nilsson, 2006). This avoids saturation of the photoreceptors and is a mechanism to maintain colour constancy (Foster, 2011; Kemp et al., 2015). Our results therefore suggest that green LEDs are discriminated based on opponent processing. In the longer wavelength range above 550 nm, yellow LEDs are presumably discriminated mainly by different stimulation of the green receptor with only low inhibitory input from a blue receptor. At darker conditions and relatively bright LED light, the green receptor might have been saturated resulting in similar signals for different wavelengths. Constant wavelength discrimination should then be possible only in the green region with distinctly overlapping receptor sensitivities resulting in different inhibitory input from a non-saturated blue receptor.

### Blue-green chromatic mechanism

The results from blue-yellow mixing experiments provide the strongest evidence for blue-green opponency (Fig. 3E,F). Small amounts of blue light decreased the preference for yellow LEDs, and thus inhibited the elicited ‘settling response’ to some extent. This reveals the presence of a blue photoreceptor with inhibitory input to an adjacent green receptor. The inverse response resembles an action spectrum of opponent inhibition and enables a first approximate estimation of the spectral location of the blue receptor (Fig. 5). These results expand the study of Stukenberg et al. (2015) which already showed that the attractiveness of green LEDs is suppressed when simultaneously combined with blue LEDs. Similarly, a blue-green chromatic mechanism was identified in the mate finding behaviour of the glow-worm *Lampyris noctiluca* also using the technique of mixing green and blue LEDs (Booth et al., 2004).

Descriptive evidence for the blue-green chromatic mechanism comes from the empirical colour choice models built from the green response and the blue inhibition experiments (Tab. 3, Fig. 7A,B). Both models explain the observed colour choice behaviour and fit well into the theory of opponent processing based on the difference of concurrent excitations of the green and blue photoreceptors. Similar models have already been shown for aphids or the pollen beetle (Döring et al., 2009; Döring et al., 2012, Döring and Röhrig). In contrast to the reported studies which were based on physiological and behavioural data, reliable physiological data on photoreceptor sensitivities were not available for *T. vaporariorum*. Therefore our flexible approach does not enable us to estimate exact positions of the photoreceptors since linear modelling based on excitation differences of several combinations of blue and green photoreceptor peak sensitivities led to well-fitting linear regressions (Tab. 3). The preference restriction that the highest receptor excitation should correspond with the LED of highest response is thereby fulfilled either in one or the other model. The position of the green receptor is limited to longer wavelengths by the preference restriction in the ‘green response model’ while the ‘blue inhibition model’ sets a limit towards shorter wavelengths. Both models follow a slightly different pattern with receptor peaks either far away from each other or close together but have a converging area in the range where receptors are close together and the restrictions are fulfilled in both models. These four combinations are 480 & 515 nm, 485 & 510 nm, 490 & 505 nm, and 495 & 500 nm (Tab. 3) which all lead to similarly shaped theoretical action spectra peaking at 554 - 556 nm (Fig. 5). While the very close combinations appear quite unlikely with regards to a reliable signal from the opponent mechanism, the more distant combinations (480 & 515, 485 & 510 nm) appear relatively realistic (Fig. 5). In comparison, the known photoreceptor sensitivities of aphids, which are also phloem-sucking herbivores show similar configurations. Receptor peaks for the green peach aphid *Myzus persicae* were determined around 490 and 530 nm and for the pea aphid *Acyrthosiphon pisum* at 518 nm, respectively (Kirchner et al., 2005; Döring et al., 2011). However, the exact positions and sensitivities of photoreceptors in the greenhouse whitefly still remain uncertain from this study, but only within a small range: The blue photoreceptor should be present with a peak around 480 - 490 nm, while a green receptor exists between 510 - 520 nm. The presence of a green receptor around 520 nm is also supported by the former ERG recording by Mellor et al. (1997). Obviously, this ERG investigation did not detect the blue receptor and measured a mixed peak of the green and blue receptor. It is unclear why the green peak was so prominent in ERG recordings but the blue photoreceptor cells may be underrepresented and contribute only a low electrophysiological input which is then strongly weighted in the nervous system.

The possible reasons for the incongruence of both models and the inaccuracies of their outcomes are diverse because they rely on simple assumptions and incalculable factors. The sensitivity functions of photoreceptors based on template formulas could slightly differ from real sensitivities for various reasons like self-screening properties or filter and screening pigments. Moreover, the calculations from photon catches to excitation values by the nonlinear transformation might not explain the reality completely. Furthermore, the relative contributions of the inputs from blue and green photoreceptors most likely differ from the assumed one-to-one ratio. Possible reasons for this could be different amounts of blue and green-sensitive photoreceptor cells in the compound eye or different weighting of the signals in the nervous system (Warrant and Nilsson, 2006; Cronin et al., 2014).

### Intensity dependence in the ‘settling response’

It could be shown that the ‘settling response’ exhibits a clear intensity dependence (Fig. 6) which is in line with findings in whiteflies and other insects (Coombe, 1981; Scherer and Kolb, 1987; Booth et al., 2004). Normally, colour vision is characterized to be independent of intensity and most studies implicate that behaviours are processed either purely chromatic or achromatic and it often remains unclear if both aspects are involved (Kelber and Osorio, 2010). But our results demonstrate that the suggested dichromatic mechanism shows both chromatic and achromatic properties, hence both colour (wavelength) and intensity are crucial in the ‘settling’ behaviour. This is an aspect which has already been implied by the colour choice model (see above) since excitation values as outcome of the opponent mechanism can theoretically be increased at the same wavelength by increasing their intensity. Our results show that within the green-yellow range of the action spectrum higher intensities can compensate for not optimally attractive wavelengths, thus colour constancy is not completely achieved. Furthermore, the sensitivity to relative intensity changes was higher in case of green LEDs compared to yellow LEDs (Fig. 6A). This represents a further clue that an interaction between receptors takes place, as these intensity dependencies would be parallel if they are based only on one receptor, following the principal of univariance (Naka and Rushton, 1966). Obviously, the intensity dependence is more distinct and stable in the green region in which the action spectrum is mainly shaped by opponent processing as compared to the yellow region where it should be primarily formed by the sensitivity of the green receptor.

Also, amongst equally coloured yellow LEDs preferences follow a brightness gradient which further demonstrates the influence of intensity on the choice behaviour in a multiple-choice setup (Fig. 6B). The interaction between the relative preferences and the ambient light intensity may be explained with photoreceptor adaptation (Fig. 6C), as has already been discussed for the wavelength choice experiments. Under bright background light conditions, the relative receptor sensitivity might be lower resulting in higher relative attractiveness of the two brightest LEDs. Under darker conditions, the relative sensitivity was probably higher resulting in a more even attractiveness of the traps.

### UV response

The moderate attraction of the greenhouse whitefly to UV radiation supposed to be a wavelength-specific behaviour involved in flight initiation, migration, and dispersal could be confirmed in our study. Apparent differences in the choice behaviour compared to the experiments in the green-yellow range corroborate that another antagonistic behaviour aside from ‘settling’ is most likely the reason for the attraction (Coombe, 1981; 1982; Stukenberg et al., 2015). One important indication was the low speed of orientation and generally low recapture rates resulting in long trial durations to achieve sufficient numbers of trapped individuals. Moreover, it could be visually observed that the orientation was not as target-oriented as the response to green since individuals tended to rest somewhere in the upper part of the cage before the traps were approached.

A relatively ambiguous wavelength dependence was determined with no significantly distinguished LEDs in the UV range below 400 nm and a high variance of the choice data (Fig. 3G). The attractiveness decreased at 400 nm but was still present in the blue range (Fig. 3H), indicating a relatively wide sensitivity. Nevertheless, the half-sided action spectrum can most likely be attributed to a uniform behaviour (Fig. 5). The observed peak of the action spectrum at 373 nm allows no final conclusion about the most attractive UV LED because we could not test high power LEDs with smaller wavelengths as they are not yet available. According to these results, it could be assumed that the observed behaviour is based only on one receptor in the UV range because no indication for a chromatic interaction with an adjacent receptor could be found. This is also supported by the colour choice model which explains the data best with a receptor peak sensitivity at 360 nm (Fig. 5, 7C). Therefore, it can be concluded that the position and peak sensitivity of the UV receptor lies between 340 and 370 nm supporting the existing study by Mellor et al. (1997) showing a UV peak around 340-350 nm in ERG recordings.

However, the conclusion that the UV receptor does not at all interact with another receptor in a behavioural context might be misleading and has to be scrutinized carefully. In a natural environment, significant intensities of UV radiation are always associated with skylight which contains all wavelengths while the light reflected from natural objects usually contains relatively small amounts of UV. It is suggested that insects could generally use a threshold-based UV-green contrast to detect skyline features and to perform landmark navigation tasks (Möller, 2002). This opponent interaction with inhibitory input from a distant green receptor would enable insects to discriminate the sky from terrestrial objects in most cases. A UV-green contrast allows a better discrimination in this context than an assumed UV-blue contrast. Therefore, it might not be the total intensity but rather the UV-green ratio in the perceived light which determines the classification into sky and object. Light with a UV ratio above a certain threshold might be classified as sky while objects with a lower UV ratio should theoretically appear as a dark silhouette.

We can assume that the UV radiation emitted by the traps in our setup competes with the UV radiation naturally entering the cage, thus skylight and trap should appear similar in this behavioural context. Theoretically, the UV traps in our setup could be perceived by the whiteflies as additional entry points for skylight which elicit an ‘open-space reaction’ as described for butterflies (Scherer and Kolb, 1987). Similarly, UV patches could be used by *T. vaporariorum* in the natural environment to find a way out of a plant canopy in order to conduct dispersal flights. But it is important to note that the solitary UV radiation emitted from LEDs in our setup is highly artificial as compared to the green-yellow LEDs which basically imitate host plants. Although not much UV radiation is transmitted through the greenhouse glass, the UV intensity measured at the release point was frequently higher than received from the traps. Only with closer distance to the traps the UV intensity became higher compared to skylight. The reason why the traps with comparably low UV intensities under such daylight conditions were attractive for whiteflies could be the mentioned UV-green ratio (Möller, 2002) which should be high due to the lack of any green light. The possibility of a UV-green contrast coincides with the antagonistic character of the behavioural pattern towards UV as compared to green (Coombe, 1981; 1982). The rationale of such a UV-green contrast mechanism to discriminate sky and object represents a convincing explanation but needs further investigations in the future.

## Conclusion and Outlook

Translated into the natural environment, *T. vaporariorum* uses the discussed chromatic mechanism to extract a colour signal to decide if the perceived object is a host plant or not. All objects with significant reflection in the green-yellow range (500-600 nm) and low reflection in the violet-blue range (400-500 nm) are potentially seen as host plants and elicit settling. Within the green-yellow range of the action spectrum the whitefly selects the brightest stimulus with the highest excitation of the opponent mechanism. The intensity dependence may be used to detect young leaves of brighter green or top leaves exposed to the sun and showing higher reflectance or transmittance than shaded ones.

This study represents the first detailed LED-based investigation on whitefly visual behaviour resulting in LED-based action spectra under natural sunlight conditions. This has profound relevance for the basic understanding of the visual mechanism of *Trialeurodes vaporariorum* and provides the basis for the improvement of visual trapping methods for monitoring and control in greenhouses. A recent study already showed the effect of modifying the reflective properties of yellow card traps (Sampson et al., 2018). This can be further specified according to the determined wavelength and intensity dependence, the photoreceptor sensitivity estimations, and the colour choice model from our study. Moreover, LEDs which enhance the attractiveness of visual traps (Stukenberg et al., 2015) can be more appropriately selected according to our results. The developed method generally provides great possibilities for future studies on the visual ecology of insects. An important notion is that we obtained comparable results regarding spectral efficiencies with multiple-choice and dual-choice experiments, thus for rapid wavelength screenings time consuming dual-choice experiments could be neglected in the future. The method could also be extended to more detailed LED mixing experiments under controlled ambient light conditions to provide a better basis for more precise modelling of photoreceptor sensitivities and interactions. Future studies should especially focus on the behaviour related to UV radiation and the underlying mechanisms.

## Acknowledgements

We gratefully acknowledge Prof. Dr. Thomas Döring for a fruitful discussion on the colour choice model, Dr. Frank Schaarschmidt for statistical advice and Dr. Christine Dieckhoff for valuable comments on language and style.

